# Potential Mechanisms Underlying Kaempferol-Promoted Osteoblast Proliferation and Osteogenic Differentiation: A Network Pharmacology and Experimental Validation Study

**DOI:** 10.1101/2025.04.02.646873

**Authors:** Huixi Wang, Zhengwei Luo, Canbin Zhao, Dongfeng Chen, Xiaolong Yan, Donghui Guan

## Abstract

**Background:** Osteoporosis (OP) manifests primarily in middle-aged and elderly individuals, representing an age-related condition characterised by diminished bone mass and alterations in bone tissue structure, potentially resulting in fractures and compromising the patient’s quality of life. The potential of kaempferol to modulate osteogenic differentiation, enhance bone metabolism, and potentially offer therapeutic benefits in OP cases is of particular interest. This study, employing a combination of network pharmacology and experimental verification, investigated the underlying mechanisms by which kaempferol stimulates the proliferation and osteogenic differentiation of mouse embryonic osteoblast precursor cells MC3T3-E1 subclone 14 through the PI3K/AKT signalling pathway. The findings provide a rational foundation for the potential of kaempferol to promote osteogenesis and enhance the treatment of OP.

**Methods:** The present study identified target genes regulated by kaempferol during osteogenesis and differentiation using network pharmacology. To this end, a protein-protein interaction (PPI) network was constructed, and Gene Ontology (GO) and Kyoto Encyclopedia of Genes and Genomes (KEGG) enrichment analyses and molecular docking were performed.The cytotoxicity of kaempferol was assessed using the CCK-8 method and the cell clone method in the cell detection section.The macroscopic regulatory mechanism of kaempferol in osteogenic differentiation was studied using alkaline phosphatase staining, activity assay, Alizarin Red staining, and calcium quantification_º_ Furthermore, real-time quantitative polymerase chain reaction (RT qPCR) and Western blot methods were employed to detect the microscopic expression differences of mRNA and protein related to the PI3K/AKT signalling pathway.The present study identified target genes regulated by kaempferol during osteogenesis and differentiation using network pharmacology. To this end, a protein-protein interaction (PPI) network was constructed, and Gene Ontology (GO) and Kyoto Encyclopedia of Genes and Genomes (KEGG) enrichment analyses and molecular docking were performed.

**Results:** A total of 203 target genes regulated by kaempferol were identified during osteogenic differentiation, with the majority of these genes being associated with biological processes related to cell proliferation and regulation.Four of these target genes act on the PI3K/AKT signalling pathway and show good interactions with kaempferol. Furthermore, kaempferol (5 μM, 10 μM) has been shown to enhance the vitality and proliferation of MC3T3-E1 Subclone 14 cells, as well as to increase alkaline phosphatase activity and calcium deposition. Furthermore, kaempferol (5 μM, 10 μM) has been observed to upregulate the mRNA expression of phosphoinositide 3-kinase (Pi3k), β-catenin, Myc proto-oncogene protein (c-Myc), and cyclin D1 in MC3T3-E1 Subclone. 14 cells, and promotes the phosphorylation of PI3K, serine/threonine protein kinase AKT (AKT1) and glycogen synthase kinase, as well as the phosphorylation of glycogen synthase kinase-3β (GSK3β) (p < 0.05), thereby upregulating the expression of β-catenin, C-MYC and CYCLIN D1 proteins and increasing the levels of p-PI3K/PI3K, p-AKT1/AKT1 and p-GSK3β/GSK3β levels, thereby promoting osteogenic differentiation of MC3T3-E1 Subclone 14.

**Conclusion:** Kaempferol has been demonstrated to have the capacity to significantly promote the osteogenic differentiation of MC3T3-E1 Subclone 14. This process is thought to be achieved by regulating the PI3K/AKT signalling pathway and affecting the expression of osteogenic-related genes. It has been shown to have a preventive and therapeutic effect on the occurrence and development of osteoporosis.

## 1. Introduction

Osteoporosis (OP) is a metabolic bone disease characterised by systemic bone loss and an imbalance between the actions of osteoblasts and osteoclasts.^[1]^_º_ The clinical manifestations of osteoporosis are characterised by a decrease in total bone mass, the destruction of bone units, and an increased propensity for fractures due to minor traumas. This condition has emerged as a significant economic and familial challenge, owing to its protracted nature, the absence of substantial therapeutic efficacy, and the high costs associated with treatment. This challenge is further compounded by the rising demographic trend of an ageing population.^[2]^,

Research has demonstrated that osteoporosis is the third most prevalent chronic disease, subsequent to hypertension and diabetes, as a consequence of inadequate consideration.^[3]^_º_The rapid growth of the elderly population has resulted in a consistent increase in the utilisation of medical services, leading to mounting pressure on the medical payment system. This necessitates urgent attention to the development of cutting-edge drugs for osteoporosis.^[4]^_º_ The primary treatment for this disease in modern medicine is Western medicine, including bisphosphonates such as alendronate sodium and RANKL inhibitors such as denosumab ^[5]^. However, it should be noted that long-term use of these drugs can result in adverse gastrointestinal reactions, dizziness, kidney toxicity, and mandibular necrosis. Moreover, while these drugs may reduce the incidence of fractures, they have been observed to increase the risk of developing tumours and cardiovascular diseases^[6]^,The consequences of the condition are manifold, impacting the patient’s treatment outcome, compliance, and long-term prognosis. Consequently, there is an urgent need to identify a pharmaceutical treatment for osteoporosis that exhibits minimal toxic side effects and optimal efficacy.

The utilisation of herbal medicine in Traditional Chinese Medicine has a long history, with numerous natural herbal products having been documented as reducing bone loss^[7]^. Among these, herbs such as Epimedium, Dipsacus, Eucommia, Pueraria, Ligustrum and Cnidium are frequently employed in combination as the primary therapeutic agents^[8]^_º_ Research has demonstrated that osteopetrolatum comprises flavonoids, which have the capacity to impede the differentiation of osteoclasts, promote the proliferation of osteoblasts, accelerate the cell cycle, and exert a favourable effect on the treatment of osteoporosis. Icariin has been shown to enhance bone and muscle strength, alleviate rheumatism, promote bone formation, inhibit bone resorption, and regulate bone reconstruction, thereby conferring a therapeutic benefit in cases of osteoporosis^[9]^_º_ As one of the primary active ingredients of osteospermum and epimedium, kaempferol has been shown to have a positive therapeutic effect on bone metabolism. Studies have demonstrated that kaempferol can mediate bone marrow mesenchymal stem cells by targeting and regulating various signal pathways, thereby regulating the differentiation, proliferation and apoptosis of osteoblasts and osteoclasts, and ultimately preventing and treating osteoporosis^[10]^. Furthermore, kaempferol has been shown to promote osteogenic differentiation of mesenchymal stem cells by regulating the PI3K/AKT signalling pathway, thereby alleviating osteoporosis in ovariectomized rats^[11]^. However, there is currently no comprehensive report on the mechanism and molecular level research on kaempferol regulating osteogenic differentiation in vitro.

Network pharmacology is an emerging discipline that provides a scientific approach and means to elucidate the mechanism of traditional Chinese medicine in the treatment of diseases. It offers a new ‘multi-target, multi-effect’ network model for this purpose. The efficacy of kaempferol in the treatment of osteoporosis has been confirmed, but its complex pharmacological effects and potential molecular mechanisms remain unclear. The present study therefore sought to explore the potential mechanism of kaempferol in the treatment of OP by regulating the PI3K/AKT signalling pathway to promote the proliferation and osteogenic differentiation of mouse embryonic osteoblast precursor cells subclone 14 (MC3T3-E1 subclone 14), using network pharmacology methods supplemented by experimental verification. The findings may provide new insights into the clinical application of OP treatment and expand related research fields.

## 2. Material and databases

### 2.1 Cell culture material

MC3T3-E1 Subclone14, as identified by STR spectrum, was tested negative for mycoplasma (Catalog number: CL-0378). The cell-specific culture medium for MC3T3-E1 Subclone 14 (Catalog number: CM-0378) was procured from Procell (Wuhan, Hubei, China). Luteolin (catalog number: S31366), crystalline purple PBS solution (catalog number: R24040), and 4% paraformaldehyde solution (catalog number: R20497) were purchased from YuanYe (Shanghai, China). The CCK-8 assay kit (catalog number: BS350B) and DMSO (catalog number: BL165B) were purchased from Biosharp (Bengbu, Anhui, China).An alkaline phosphatase staining kit (catalog number: G1480) and a total RNA extraction kit (catalog number: R1200) were purchased from Solarbio (Beijing, China). The xylene red staining solution (catalog number: G1038) was procured from Servicebio (Wuhan, Hubei, China).A CO2 incubator (catalog number: BPN-80CH) was purchased from Yiheng Instruments (Shanghai, China), and a laminar flow cabinet (catalog number: SW-CJ-1FD) was purchased from Suzhou Clean Air Technology Co., Ltd. (Suzhou, Jiangsu, China).

### 2.2 Databases

## 3. Method

### 3.1 Network pharmacology

#### 3.1.1 Screening of kaempferol target genes

Obtain the two-dimensional structure SDF file of kaempferol from the PubChem website and import it into the SwissTargetPrediction and PharmMapper drug target gene prediction tools. Export and merge the predicted target gene files to remove duplicate values to obtain the target genes affected by kaempferol.

#### 3.1.2 Select osteoblast differentiation targets and target genes

Use the Genecards database tool to search for genes related to bone formation using the keyword “osteoporosis”.

#### 3.1.3 Venn diagram of drug target genes and osteogenic differentiation-related genes

The kaempferol-affected target genes obtained in section 3.1.1 were imported into the OmicShare Tools website together with the osteogenic differentiation-related genes obtained in section 3.1.2. A Venn analysis was performed to determine the intersection of these two sets of osteogenic differentiation-related target genes representing kaempferol regulation.

#### 3.1.4 Constructprotein-protein interaction (PPI) network

The STRING database tool was used to create a PPI network using the cross-target genes obtained in 3.1.3. After exporting the protein-protein interaction network diagram, topological calculations and optimization were performed using Cytoscape 3.9.1 (UC San Diego, San Diego, CA, USA). According to the results, the core PPI network was re-screened based on the degree value (>20).

#### 3.1.5 GO enrichment and KEGG pathway enrichment analysis

The cross-target genes obtained by g:Profiler in 3.1.3 were analyzed using the dynamic GO and KEGG analysis tools. The results of GO enrichment analysis (biological processes,cellular components, and molecular functions) and KEGG pathway enrichment analysis were obtained, and bubble and bar charts were generated.

#### 3.1.6 Molecular docking verification

Proteins involved in the PI3K/AKT/GSK-3β pathway were selected from the PPI network. Molecular docking of these proteins and kaempferol was performed using Auto Dock 4.2.6 software (The Scripps Research Institute, San Diego, CA, USA). Finally, the molecular docking images were visualized and analyzed using Pymol 2.4.0 software (University of California, Oakland, CA, USA).

### 3.2 Cell experiment

#### 3.2.1 CCK-8 experiment

#### 3.2.2 Cell cloning experiment

MC3T3-E1 subclone14 cells in logarithmic growth phase were seeded at a density of 5 × 103 cells/well in a 96-well plate. The cells were treated with different concentrations of kaempferol (0 μM, 5 μM, 10 μM, 25 μM, 50 μM), and 0.1% DMSO was added as solvent to each group of kaempferol (to eliminate experimental error, the 0 μM kaempferol group contains only 0.1% DMSO). After 24, 48, and 72 hours of incubation, CCK-8 detection was performed using the prepared CCK8 reagent, followed by incubation in the dark at 37°C for 1.5 hours. Data were recorded by measuring the absorbance at a wavelength of 450 nm.

Measurement of alkaline phosphatase activity and quantification of alizarin red staining Alkaline phosphatase activity assay: MC3T3-E1 subclone 14 cells were seeded at a density of 2 × 105 cells per well in a six-well culture plate. The cells were then cultured in MC3T3-E1 osteogenic differentiation induction medium containing different concentrations of kaempferol for 7 days. Alkaline phosphatase activity was measured using the BCIP/NBT alkaline phosphatase staining kit. Xylenol Red staining and quantification: MC3T3-E1 subclone 14 cells were cultured in MC3T3-E1 osteogenic differentiation induction medium containing different concentrations of kaempferol for 14 days. Alizarin red staining was performed, and the stained calcified nodules were dissolved in 10% hexadecylpyridinium chloride (Sigma-Aldrich) at room temperature. Absorbance was measured at a wavelength of 562 nm.

Real-time quantitative polymerase chain reaction (RT-qPCR) detects the expression of cellular mRNA MC3T3-E1 subclone 14 cells were seeded at a density of 1×105/well in a six-well plate and cultured in osteogenic differentiation induction medium containing different concentrations of kaempferol for 7 days. After induction, total RNA was extracted by the TRIzol method. Reverse transcription and cDNA amplification were performed, and the results were calculated using the 2-ΔΔCt method. The primer sequences were synthesized by Beijing Ding Guo Changsheng Biotechnology Co, Ltd. The specific sequences are listed in Table 2.

**TABLE 1.** Database related information.

| Databases (website) | Website and country | Databases (website) | Website and country |
| --- | --- | --- | --- |
| PubChem | https://pubchem.ncbi.nlm.nih.gov ( China ) | SwissTargetPrediction | www.swisstargetprediction.ch ( Switzerland ) |
| PharmMapper | http://www.lilab-ecust.cn/pharmmapper/ ( China ) | Genecards | https://www.genecards.org/ (Israel) |
| g:Profiler | https://biit.cs.ut.ee/gprofiler/gost (America) | String | https://www.string-db.org ( Switzerland ) |
| Omicshare | https://www.omicshare.com/ ( China ) | PDB | https://www.rcsb.org ( America ) |
| Huayuan | https://www.chemsrc.com/casindex/ ( China ) | String | https://cn.string-db.org |

**TABLE 2.**
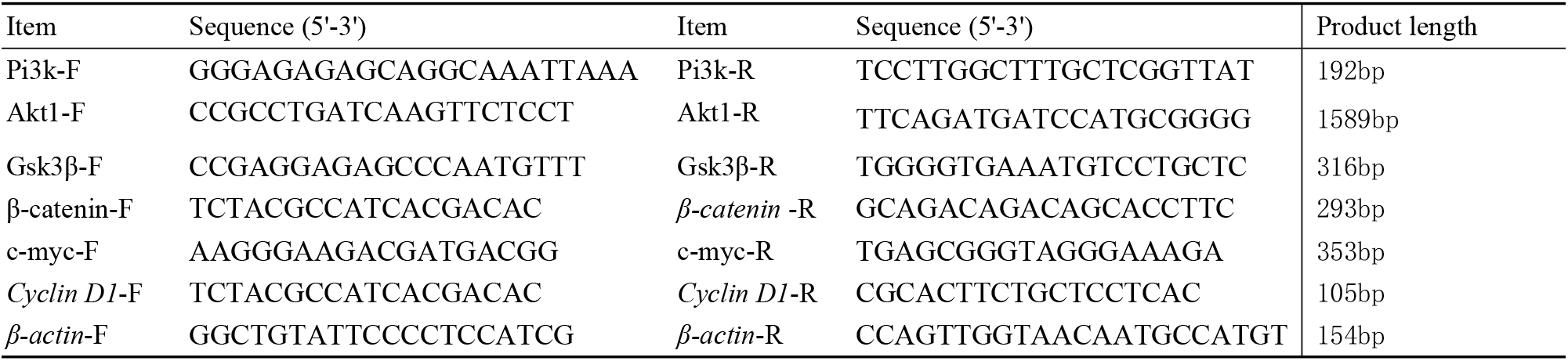
Primers for qRT-PCR detection.

#### 3.2.5 Western blot analysis of protein expression in cells

The method of drug administration was the same as that used for RT-qPCR. After drug treatment, total cellular proteins were extracted and protein concentrations were determined. Subsequently, 25 μg of protein sample per well was loaded for SDS-PAGE. Proteins were then transferred, blocked and incubated with primary antibodies overnight. The primary antibodies used were PI3K, p-PI3K, AKT1, p-AKT1, GSK3β, p-GSK3β, β-catenin, c-Myc, cyclin D1 and β-actin. After incubation with secondary antibodies for 1 hour at room temperature and washing of membranes with TBST, chemiluminescence was performed. Protein expression levels were normalised to β-actin (histone H3) standards. Band analysis was performed using Gerpol32 4.0 software (Media Cybernetics, Bethesda, MD, USA). Information on the WB antibodies is given in Table 3.

**TABLE 3.**
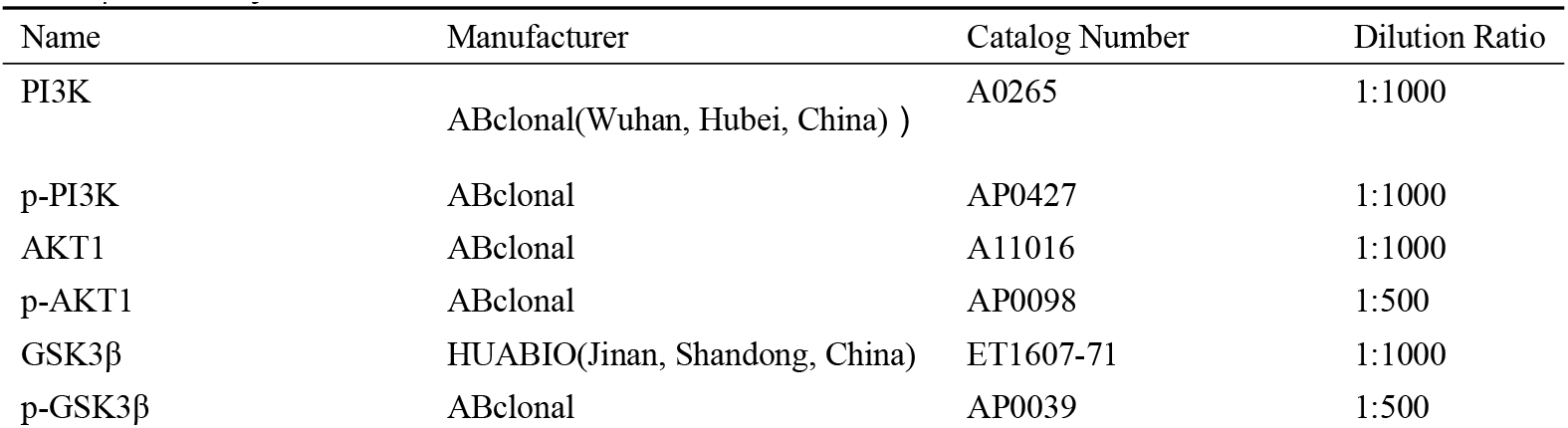
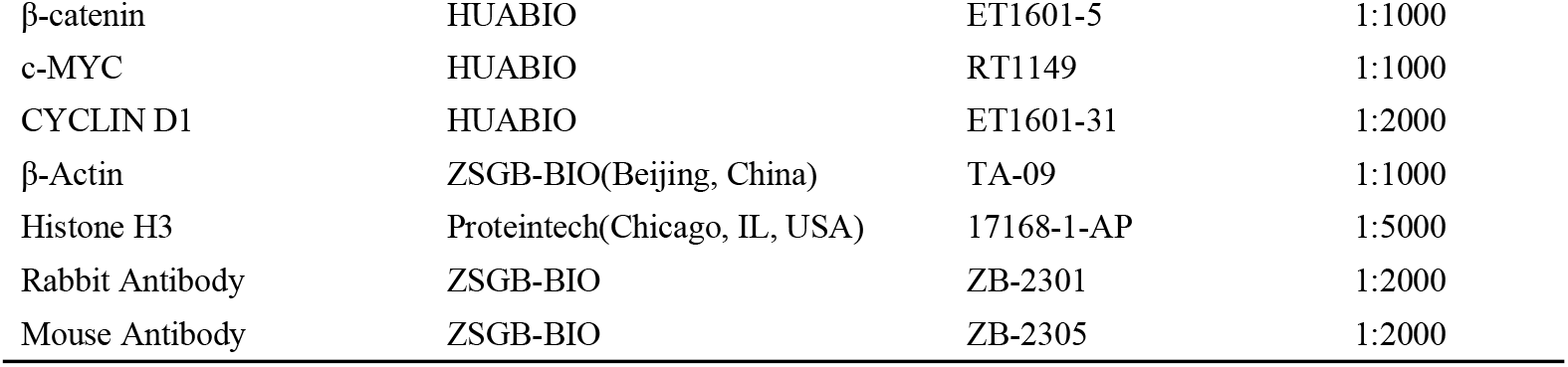
Antibody information of Western Blot.

#### 3.2.6 Immunofluorescence Detection of Nuclear Translocation of β-catenin

Cells were seeded onto culture dishes pre-prepared with treated coverslips. When the cells reached confluence, the coverslips were removed. The cells were fixed with 4% paraformaldehyde for 20 minutes. Then 300 µl of 0.3% Triton X-100 was added to each well and incubated for 20 min at room temperature. An appropriate amount of normal goat serum working solution was added to the coverslips and incubated at 37°C for 1 hour. After the addition of the primary antibody, the cells were incubated at 37°C for 2 hours.

DAPI staining solution was added and incubated for 10 minutes, followed by washing off excess DAPI. The coverslips in the wells were gently removed with tweezers and placed on glass slides. An anti-fluorescence quenching reagent was added and a coverslip was placed on top. Images were captured using a fluorescence microscope. Information on the immunofluorescence antibodies is given in Table 4.

**TABLE 4.** Antibody information of Western Blot.

| antibody information | Manufacturer | Catalog Number |
| --- | --- | --- |
| β-catenin | HUABIO | ET1601-5 |
| Cy3-labeled goat anti-rabbit IgG | Servicebio | GB21303 |

### 3.3 Statistical processing

Experimental results are expressed as mean ± standard deviation and compared between groups using one-way analysis of variance. Differences between groups were compared by t-test. Statistical analysis and graphing were performed using SPSS 21 (IBM Corp, Armonk, NY, USA) and GraphPad Prism 9 software. A p-value less than 0.05 was considered statistically significant. Each experiment was replicated three times to ensure reliability.

## 4 Result

### 4.1 Results of network pharmacology analysis

#### 4.1.1 Screening for kaempferol and osteogenic differentiation targets

The two-dimensional structure of kaempferol was obtained from the PubChem website (see Figure 1-A). The SwissTargetPrediction and PharmMapper target prediction tools identified 358 kaempferol target genes. A total of 5936 target genes related to osteogenic differentiation were retrieved from the GeneCards database.

**Figure 1:**
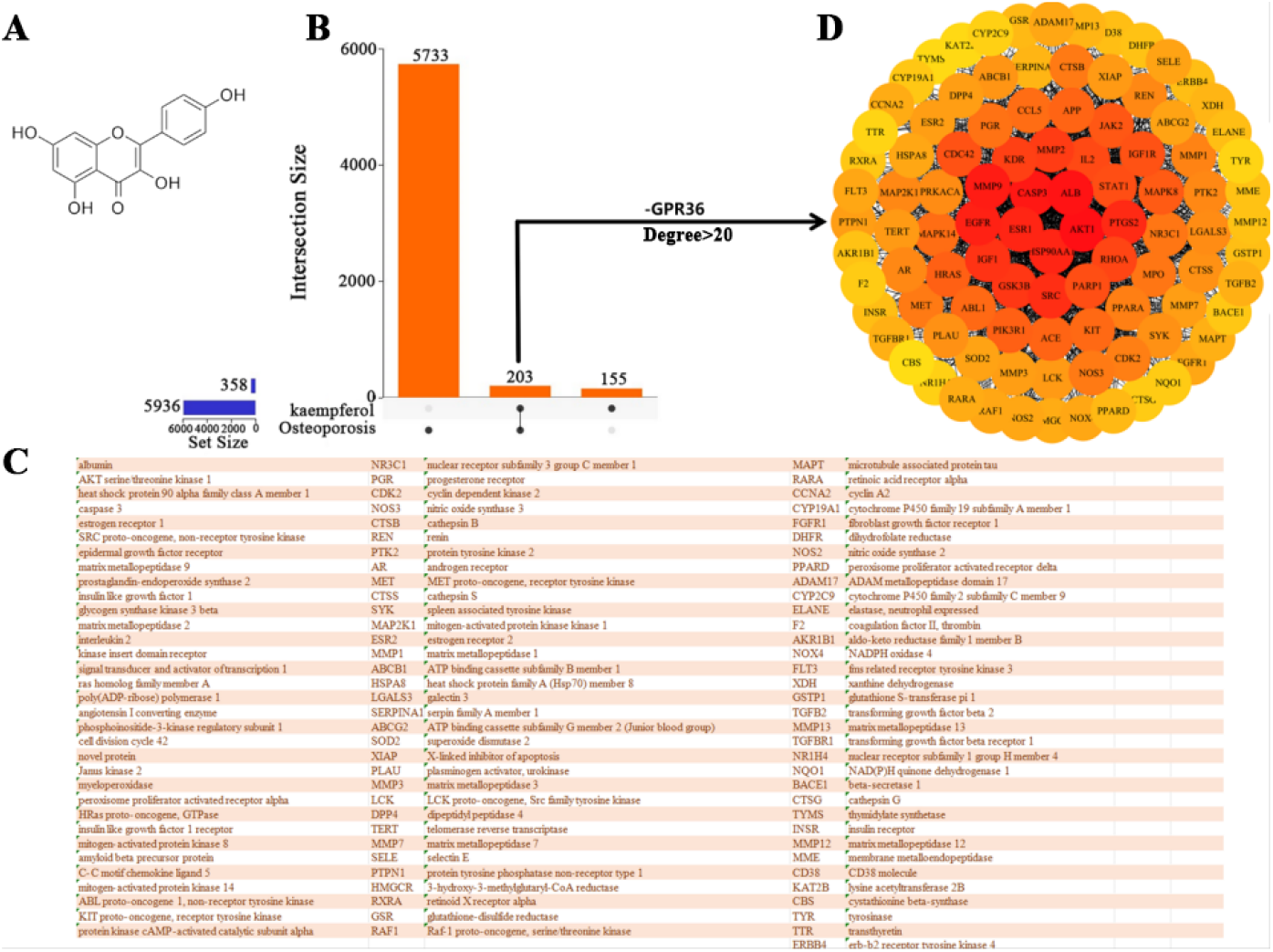
**A:**The two-dimensional chemical structure of kaempferol was obtained through network pharmacology. **B**: Venn diagram and list of kaempferol target genes and osteogenic differentiation-related genes. **C**: List of 99 genes screened by degree>20. **D**: Protein-protein interaction (PPI) core network constructed using the intersecting genes.

#### 4.1.2 Constructing a Venn diagram of drugs and disease targets

A Venn analysis was performed on 5,936 osteogenic differentiation-related genes and 358 kaempferol target genes using the Omicshare website tool. The analysis revealed that kaempferol may potentially impact 203 genes involved in osteogenic differentiation (see Figure 1-B).

#### 4.1.3 Constructingprotein-proteininteraction (PPI) networks

The STRING database was utilised to construct a protein-protein interaction (PPI) network comprising 203 genes that have the potential to be influenced by kaempferol during the process of osteogenic differentiation. With the exception of the G protein-coupled receptor 36(GPR369)gene, which was not involved in the network construction, all other genes contributed to the formation of the PPI network. A total of 99 target genes with a degree value greater than 20 were selected (see Figure 1-C), and a core PPI network was subsequently constructed (see Figure 1-D).

#### 4.1.4 GO function enrichment analysis

As illustrated in Figure 2, the results of the GO functional enrichment analysis indicate that the biological processes encompass cell proliferation, regulation of cell proliferation, and response to chemical substances. The cellular components include cytoplasm, cell membrane, and protein kinase complex. The molecular functions include small molecule binding, protein kinase activity, and anion binding.

**Figure 2:**
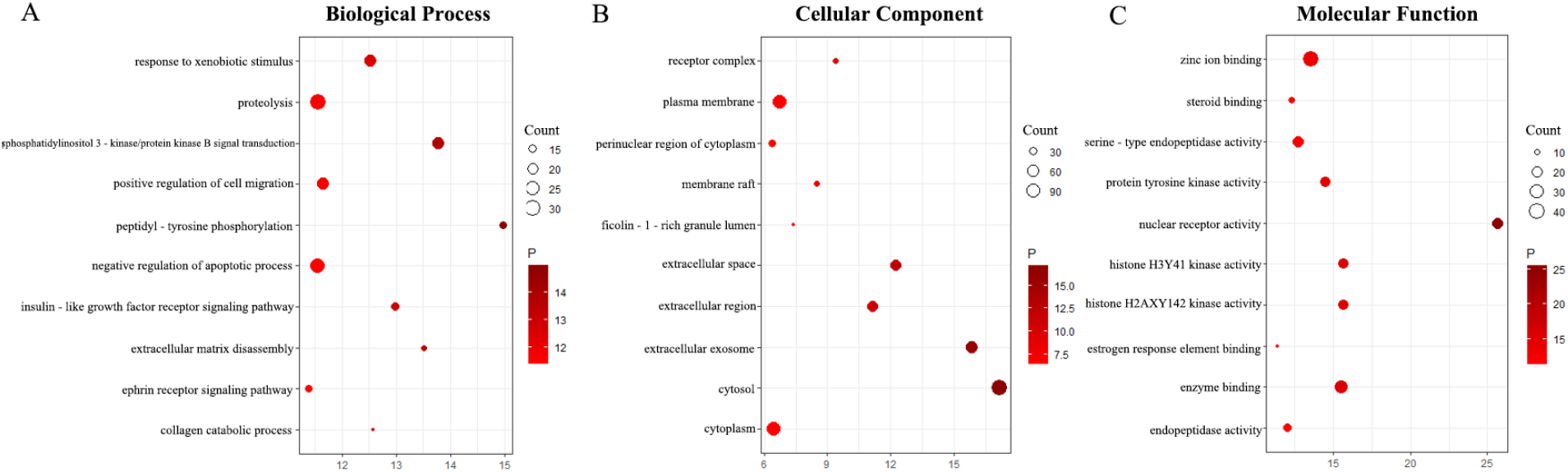
GO Bubble chart of enrichment analysis results.**A**: Biological processes. **B**: Cellular components. **C**: Molecular functions.

#### 4.1.5 KEGG pathway enrichment analysis and molecular docking

KEGG pathway enrichment analysis revealed that the genes regulated by kaempferol during osteogenic differentiation were predominantly implicated in cancer and Ras signaling pathways (see Figures 3-B and C).The PI3K/AKT signaling pathway ranked 19th in terms of importance, and within this pathway, AKT1, PI3KR1, PI3KCG, and GSK3β play a regulatory role. Molecular docking with these genes using kaempferol revealed the formation of hydrogen bonds with AKT1, PI3KR1, PI3KCG, and GSK3β proteins, with a docking energy of less than -4.25 kcal/mol ^[11]^. This observation suggests that kaempferol binds to these proteins favorably (see Figure 3-A).

**Figure 3:**
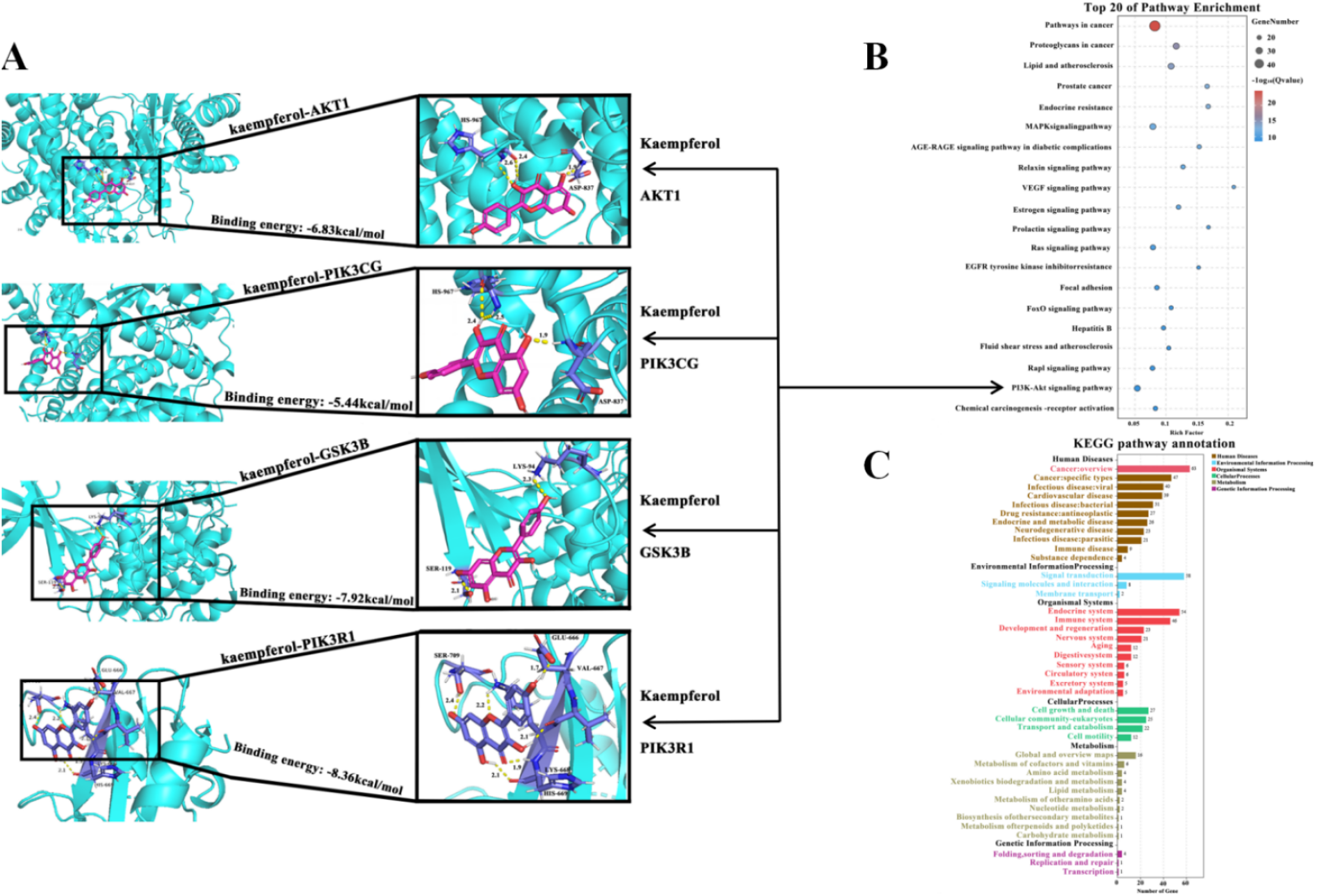
Results of KEGG pathway enrichment analysis and visualization of the docking of kaempferol with PI3K/AKT signaling pathway-related proteins._º_A: Visualization of the protein docking between kaempferol and AKT1, PI3KR1, PI3KCG, and GSK3β (the figure shows the binding site, bond length, and docking energy).B: KEGG pathway enrichment analysis bubble chart.The bar chart analysis of the number of genes involved in various biological pathways in KEGG pathways is also presented.

### 4.2 Experimental verification results

#### 4.2.1 CCK-8 cell viability assay

In comparison with the 0 μM kaempferol group, the cell viability in the 5 μM and 10 μM kaempferol groups exhibited a significant increase after 24 hours (*p* < 0.05). However, no substantial change in cell viability was observed in the 25 μM kaempferol group. Conversely, a significant decrease in cell viability was detected in the 50 μM kaempferol group (*p* < 0.05). After 48 hours of treatment, cell viability was significantly higher in the 5 μM and 10 μM shanaiol groups (*p* < 0.05), and significantly lower in the 25 μM and 50 μM shanaiol groups (*p* < 0.05). After 72 hours of treatment, cell viability increased significantly in the 5 μM kaempferol group (*p* < 0.05), while no significant change was observed in the 10 μM kaempferol group. However, a significant decrease was noted in the 25 μM and 50 μM kaempferol groups (*p* < 0.05). These findings suggest that 5 μM and 10 μM kaempferol do not demonstrate significant toxicity and possess the potential to enhance cell viability (see Figure 4).

**Figure 4:**
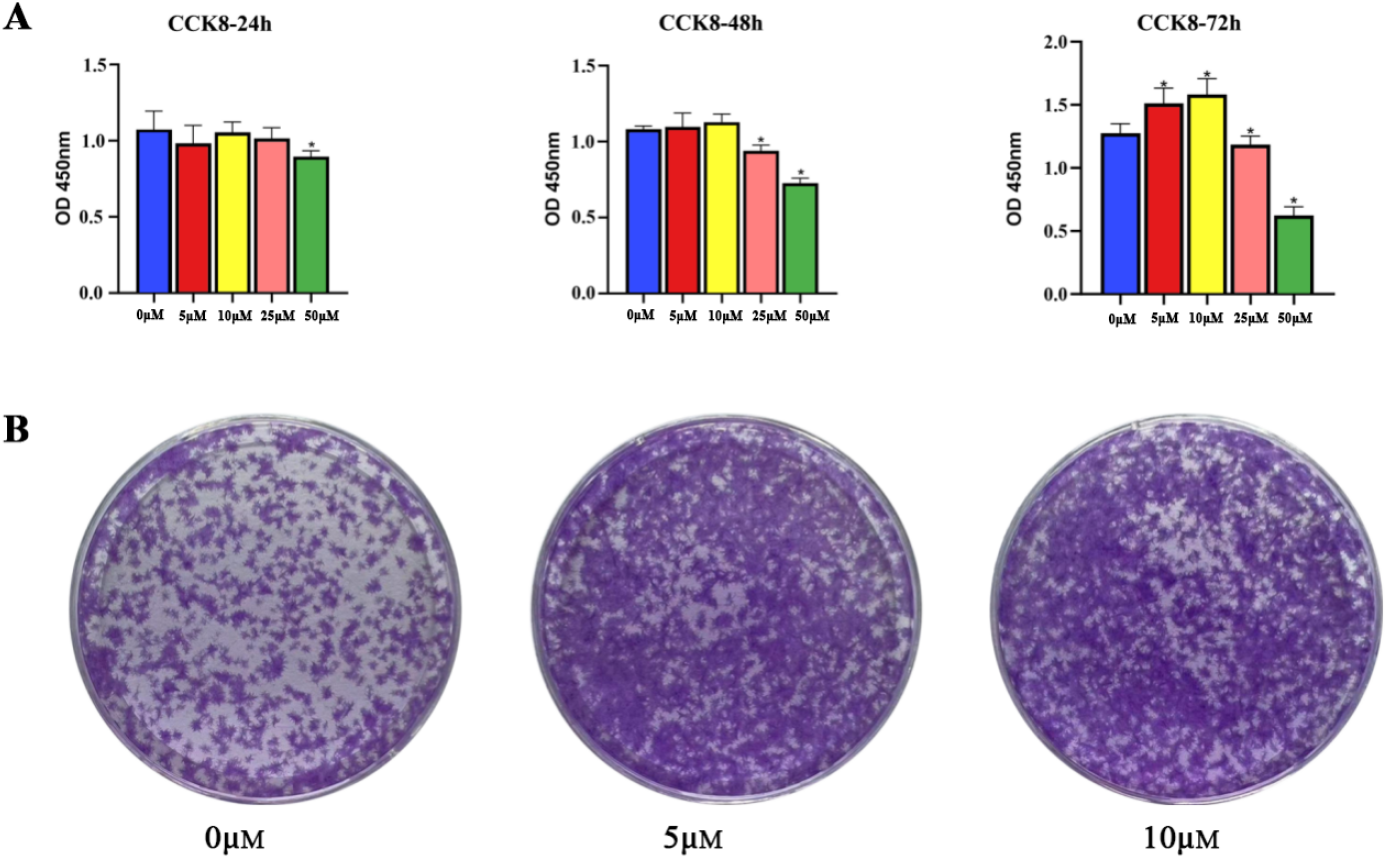
The following is a presentation of the CCK-8 and cell clone experiment results.**A:** Twenty-four, forty-eight, and seventy-two hour CCK-8 data bar charts.**B:** Results of cell clone formation experiments after stimulation with different concentrations of kaempferol in culture dishes.(*: *p* < 0.05)

#### 4.2.2 Cell cloning assay

As illustrated in Figure 4-B, the CCK-8 assay revealed that 5 μM and 10 μM kaempferol concentrations promoted cell activity. Utilizing these concentrations, a cell clone formation assay was conducted to investigate the impact of kaempferol on the in vitro proliferation of MC3T3-E1 Subclone 14 cells. A comparison of the 0 μM shikimic acid group with the 5 μM and 10 μM shikimic acid groups revealed enhanced cell proliferation, suggesting that 5 μM and 10 μM shikimic acid can promote cell proliferation.

#### 4.2.3 Alkaline phosphatase (ALP) staining and activity determination

As illustrated in Figure 5, the ALP staining results demonstrated a deeper staining intensity in the 5 μM and 10 μM kaempferol groups compared to the control group. Furthermore, the ALP activity assay results revealed a significant increase in ALP activity in the 5 μM and 10 μM kaempferol groups (p < 0.05). These findings suggest that 5 μM and 10 μM kaempferol can enhance the alkaline phosphatase activity of MC3T3-E1 Subclone 14 cells, thereby promoting osteogenic differentiation.

**Figure 5:**
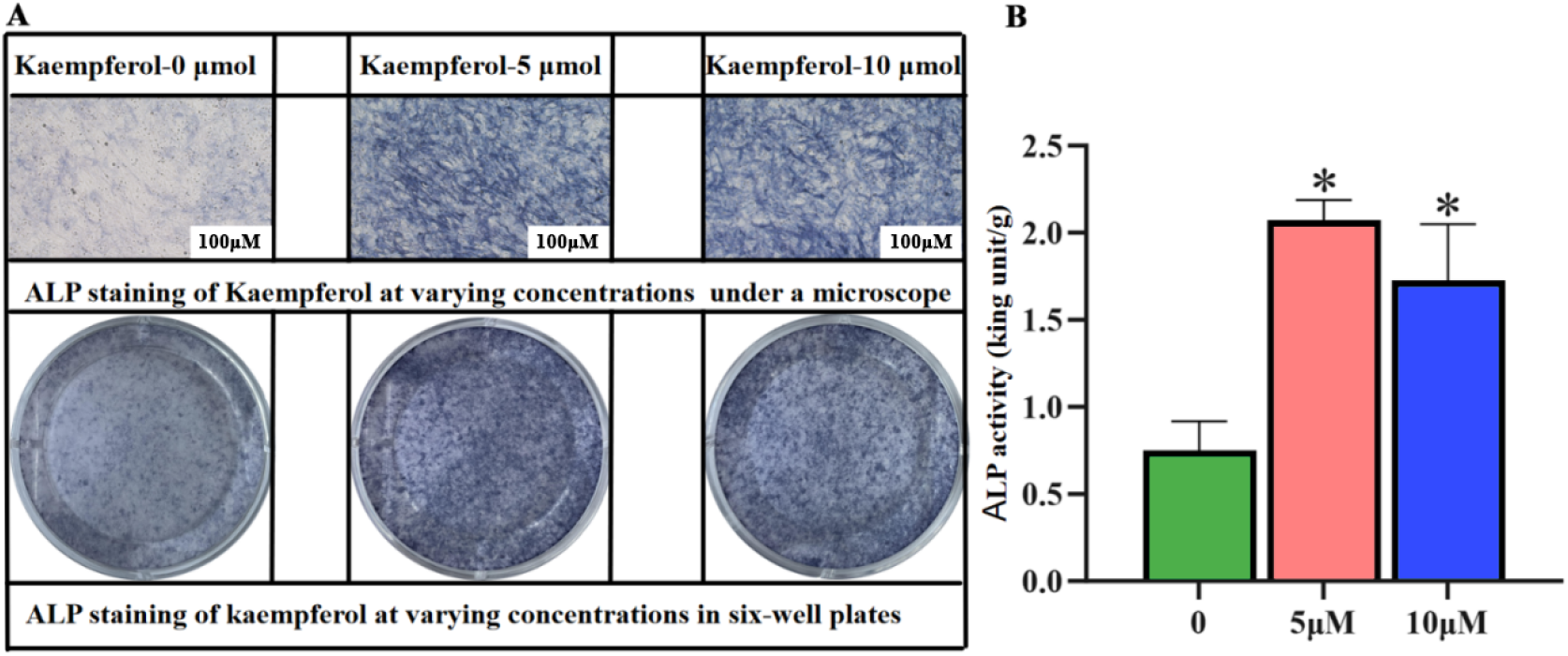
The following results are presented in relation to the staining and activity measurement of ALP.**A**: Micrographs and corresponding appearance (100×) of ALP staining in a six-well plate under the stimulation of kaempferol at different concentrations.**B**: Bar chart of ALP activity measurement results (*: *p* < 0.05).

#### 4.2.4 Alizarin red staining and quantitative analysis

As illustrated in Figure 6, alizarin red staining revealed a conspicuous enhancement in staining intensity in the 5 μM and 10 μM kaempferol groups when compared to the control group. This observation was accompanied by a notable increase in calcium deposition.Quantitative analysis of alizarin red staining indicated a significant surge in alizarin red content in the 5 μM and 10 μM kaempferol groups (*p* < 0.05). This finding suggests that 5 μM and 10 μM kaempferol can enhance intracellular calcium deposition, thereby promoting the in vitro osteogenic differentiation of MC3T3-E1 Subclone 14 cells.

**Figure 6:**
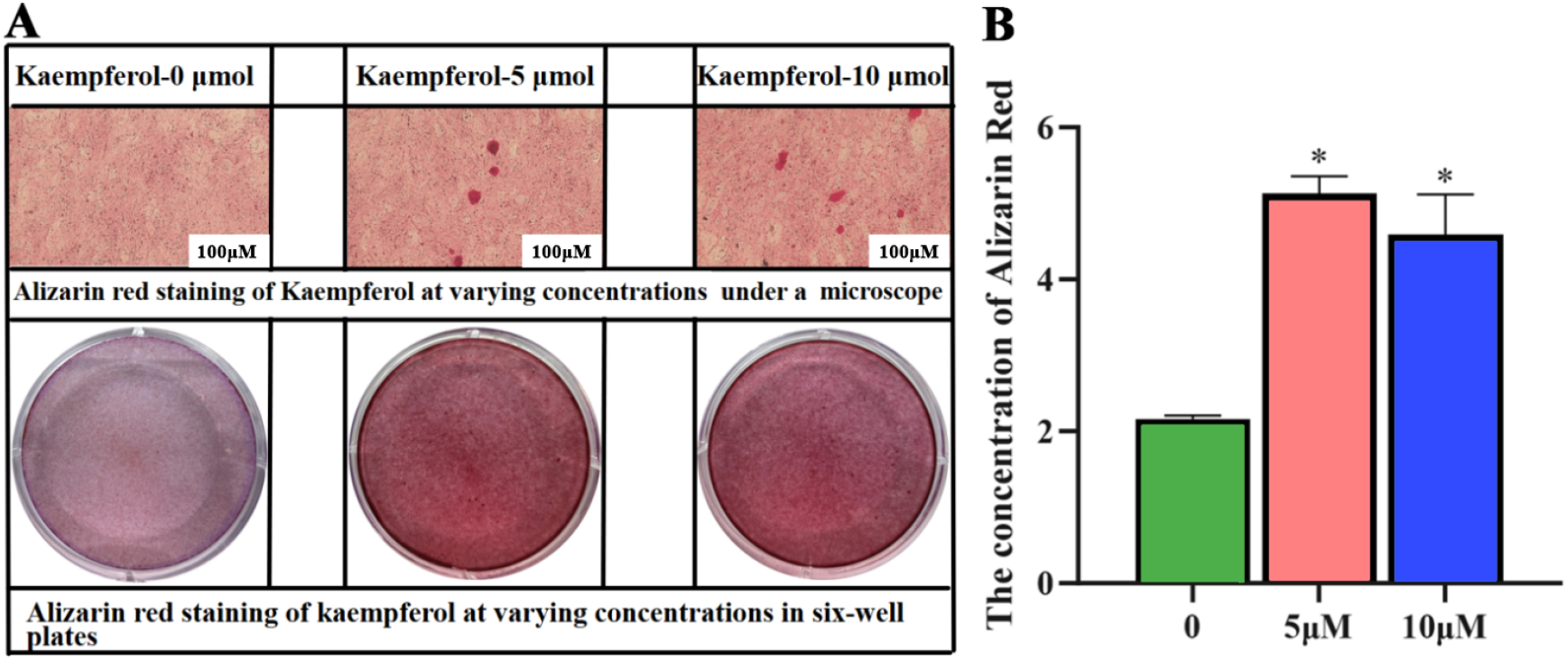
Quantitative analysis of alizarin red staining and calcium deposition. **A:** The performance of alizarin red staining at different concentrations in a six-well plate and the corresponding microscopic images (100×). **B:** Bar chart of quantitative results of alizarin red staining (*: *p* < 0.05).

#### 4.2.5 Analysis of cellular mRNA expression by RT-qPCR

As illustrated in Figure 7, the RT-qPCR outcomes revealed that, in comparison with the control group, the mRNA expression of β -catenin, c-Myc, Cyclin D1, and Pi3k was significantly elevated in the 5 μM and 10 μM kaempferol groups.Conversely, no substantial alterations were observed in Akt1 and Gsk3β mRNA expression in the 5 μM and 10 μM kaempferol groups. These results suggest that 5 μM kaempferol can promote Pi3k mRNA expression, while 5 μM and 10 μM kaempferol can upregulate β-catenin, c-Myc, and Cyclin D1 mRNA expression, which may contribute to osteogenesis.

**Figure 7:**
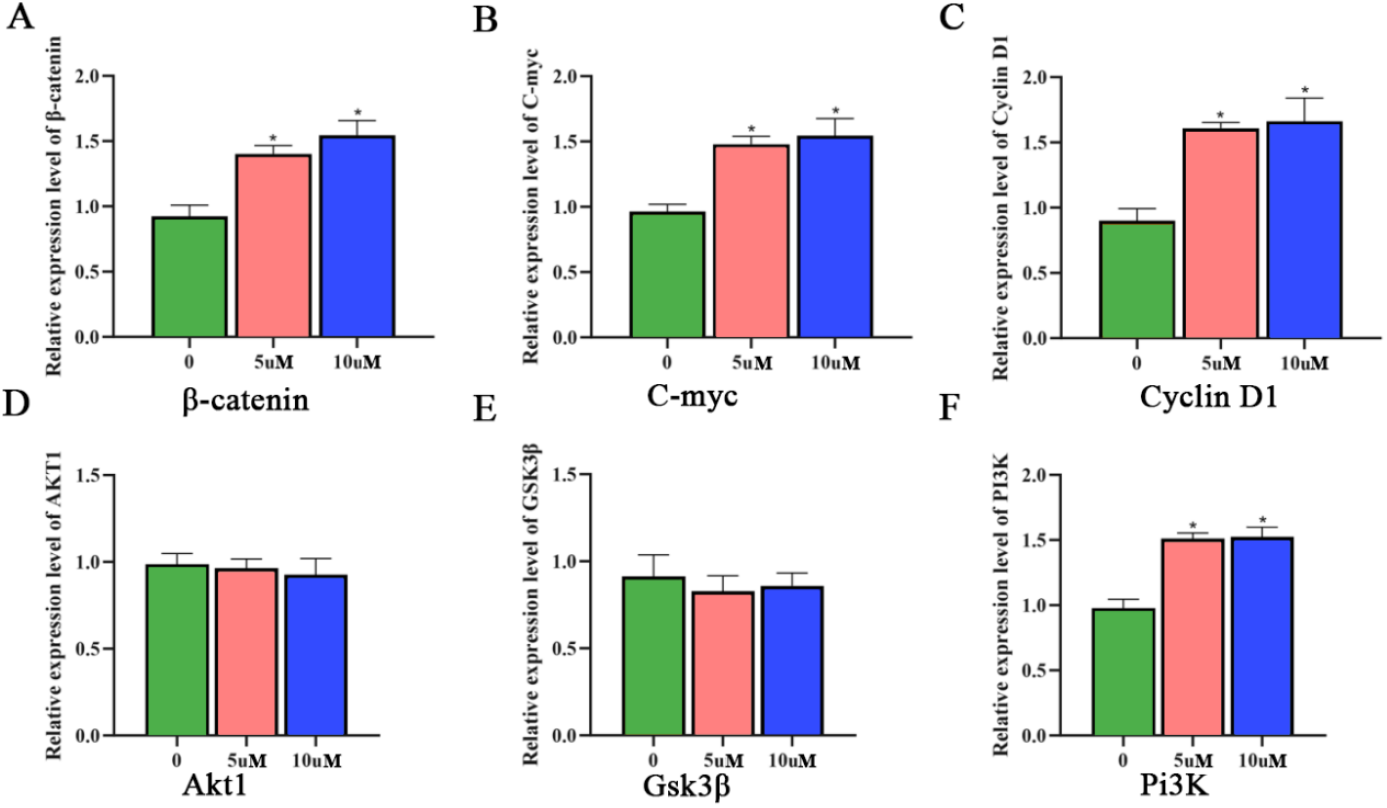
Intracellular mRNA expression was analyzed, revealing the following results:**A**: β-catenin **B**: c-Myc **C**: Cyclin D1 **D**: Akt1 **E**: Gsk3β **F**: Pi3k (*: *p* < 0.05).

#### 4.2.6 Expression and analysis of key proteins in the PI3K/AKT signaling pathway

As illustrated in Figure 8, the ratios of p-PI3K/PI3K, p-AKT1/AKT1, and p-GSK3β/GSK3β in the 5 μM and 10 μM kaempferol groups were significantly elevated compared to the control group (*p* < 0.05). Concurrently, the expression levels of c-MYC and CYCLIN D1 were found to be significantly elevated (*p* < 0.05).These findings suggest that 5 μM and 10 μM kaempferol can effectively activate the PI3K/AKT signaling pathway by phosphorylating PI3K and AKT1. This, in turn, results in the phosphorylation of GSK3β, which, in turn, promotes the nuclear accumulation of β-catenin, subsequently initiating downstream gene transcription and enhancing cell proliferation.

**Figure 8:**
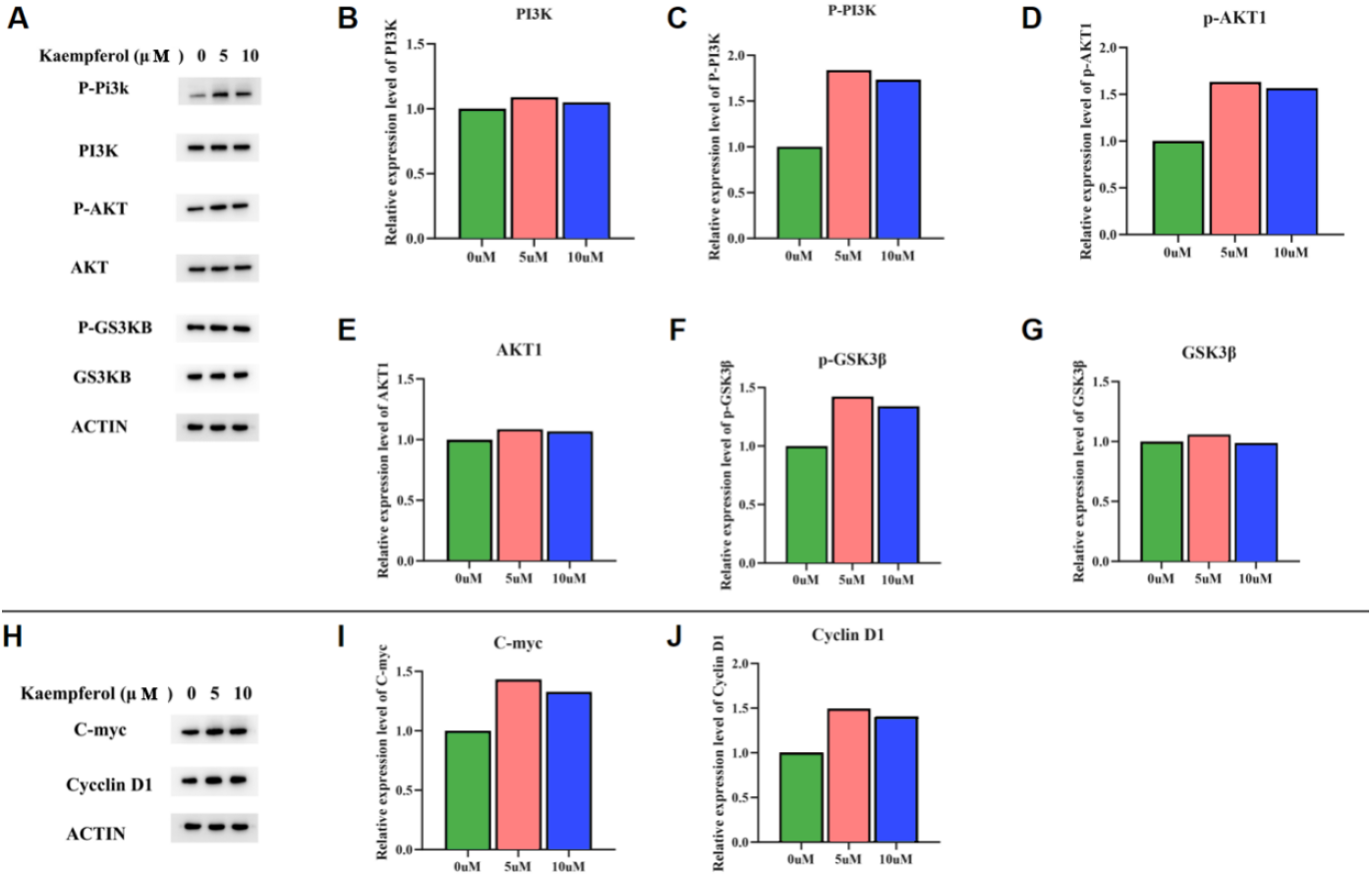
The present study investigates the expression levels of key proteins in the PI3K/AKT signaling pathway and the subsequent data analysis. **A:**the expression levels of the proteins PI3K, AKT1, GSK3β, and the phosphorylated proteins p-PI3K, p-AKT1, p-GSK3 β, and their corresponding bands under treatment with different concentrations of kaempferol. **B-G:** The numerical trends of protein expression levels under treatment with different concentrations of kaempferol (**B, C:** p-PI3K/PI3K, **D, E:** p-AKT1/AKT1, **F, G:** p-GSK3 β/GSK3β), **H:** the expression levels of c-MYC and CYCLIN D1 downstream of the PI3K/AKT signaling pathway and their corresponding bands under treatment with different concentrations of kaempferol, **I:** c-MYC, **J:** CYCLIN D1. (*: *p* < 0.05).

#### 4.2.7 Western blot and immunofluorescence analysis of the core protein β-catenin

As illustrated in Figure 9, the results of the western blot analysis revealed that there was no substantial alteration in the cytoplasmic β-catenin levels of MC3T3-E1 Subclone 14 cells treated with 5 μM and 10 μM kaempferol, in comparison to the 0 μM kaempferol group. However, a significant increase in nuclear β-catenin was observed (p < 0.05), accompanied by a substantial increase in total β-catenin expression (p < 0.05). The immunofluorescence results demonstrated that, in MC3T3-E1 Subclone 14 cells treated with 5 μM and 10 μM shikimic acid, there was a significant nuclear translocation of β-catenin when compared with the 0 μM shikimic acid group. The findings from both Western blot and immunofluorescence analyses demonstrated that 5 μM and 10 μM kaempferol could enhance the expression of β-catenin protein in MC3T3-E1 Subclone 14 cells and promote the nuclear translocation of β-catenin.

**Figure 9:**
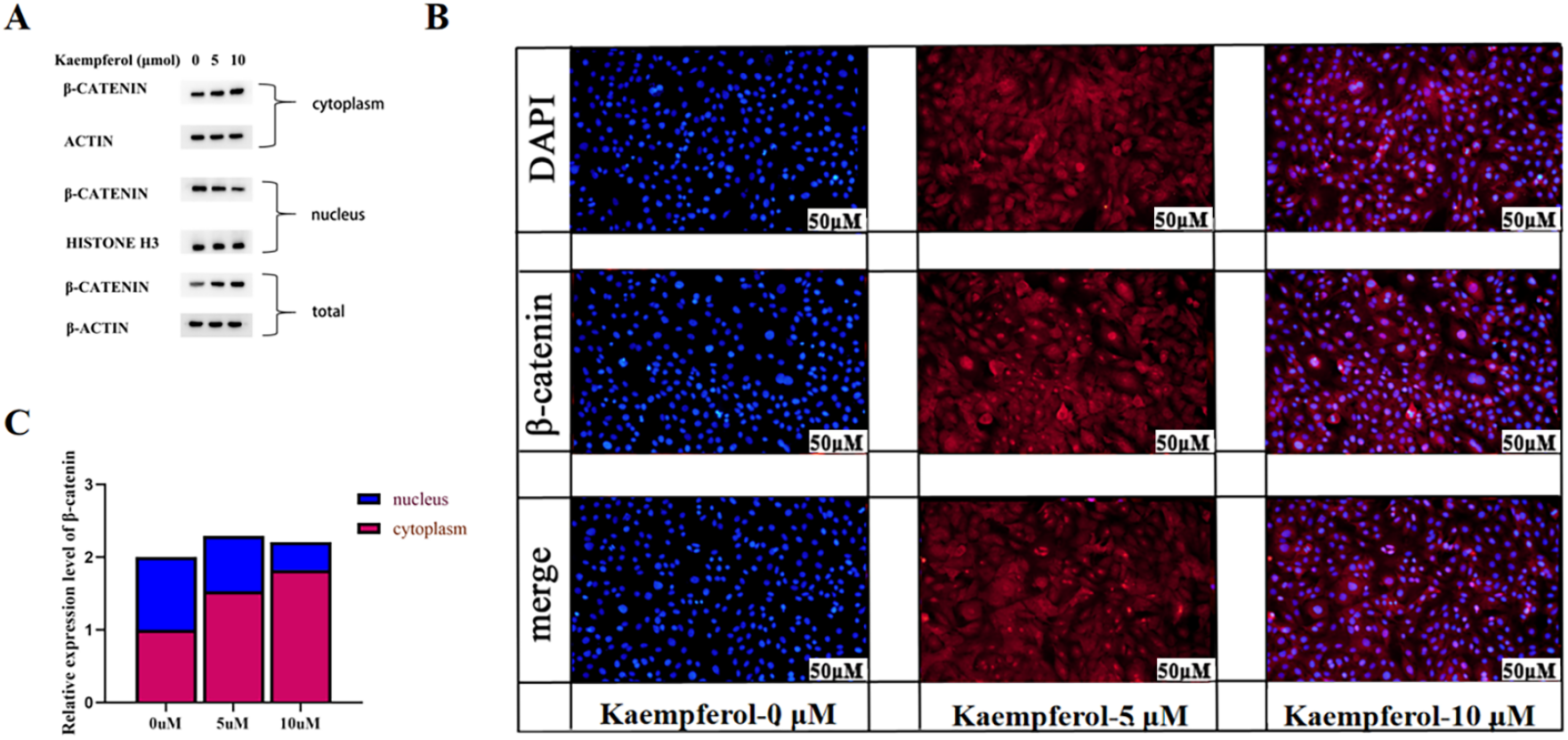
Expression and nuclear translocation of the core protein β-catenin. A: Expression levels of total β-catenin and β-catenin in the cytoplasm and nucleus and their corresponding bands under treatment with different concentrations of kaempferol. B: Nuclear translocation of the core protein β-catenin under different concentrations of Luteolin treatment. C: Expression trends of β-catenin and β-catenin in the cytoplasm and nucleus under treatment with different concentrations of kaempferol. (*: *p* < 0.05).

## 5 Discussion

Osteoporosis patients pose a serious challenge to the safety of life and property in daily life due to their pathological characteristics, such as body pain and tendency to fracture with minor force^[12]^.Osteoporosis has emerged as a social problem that is increasingly impacting families and the nation. This is primarily due to the prolonged nature of the treatment cycle, the absence of substantial treatment effects, and the exorbitant costs associated with treatment^[13]^.Research has demonstrated that, due to a myriad of factors present in daily life, individuals frequently possess a hazy understanding of its prevention and treatment. Moreover, the prevalence of osteoporosis among chronic diseases continues to escalate. While pharmaceutical interventions, namely RANKL inhibitors, calcitonin, and selective estrogen receptor modulators, which constitute the primary treatment modality, have been demonstrated to ameliorate symptoms, they frequently result in significant adverse effects and are ineffective in reducing the risk of fracture^[14-15]^.Kaempferol has been shown to mediate the function of bone marrow mesenchymal stem cells by targeting and regulating multiple signal pathways. This, in turn, has been demonstrated to regulate the differentiation, proliferation, and apoptosis of osteoblasts and osteoclasts. Consequently, kaempferol has the potential to prevent and treat osteoporosis. This finding suggests a promising avenue for the treatment of osteoporosis^[16-17]^.The PI3K/AKT signaling pathway plays a vital role in regulating osteogenic differentiation and bone formation. Research has demonstrated that the PI3K/AKT pathway modulates the proliferation, migration, and invasion of osteoblasts ^[18]^ and is intimately associated with the osteogenic differentiation of bone marrow mesenchymal stem cells ^[19]^. Inhibition of the PI3K/Akt pathway has been shown to reduce the expression of osteogenic-related genes, including ALP, OCN, Osterix, and Runx2, thereby impeding osteoblast differentiation ^[20-21]^. This study identified a network of genes, potentially influenced by kaempferol during osteogenic differentiation, including four target genes that regulate the PI3K/AKT pathway: PI3KR1, PI3KCG, AKT1, and GSK3β, which are all critical components of the PI3K/AKT signaling pathway.Research has demonstrated that phosphorylated AKT can activate the PI3K/AKT signaling pathway and phosphorylate GSK3β, thereby regulating the Wnt/β-catenin pathway and affecting bone metabolism ^[22]^. Following phosphorylation, GSK3β becomes inactive, resulting in the disassembly of the degradation complex within the Wnt/β-catenin pathway. This, in turn, results in the accumulation of β-catenin within the cytoplasm and its subsequent entry into the nucleus, where it initiates downstream gene transcription ^[23-24]^. Cyclin D1 and c-Myc, downstream target genes of the Wnt/β-catenin signaling pathway, play an instrumental role in the osteogenic process. The upregulation of cyclin D1 and c-Myc expression has been shown to accelerate osteogenic differentiation ^[25-26]^. The present study found that 5 μM and 10 μM kaempferol can upregulate the mRNA expression of β-catenin, c-Myc, and Cyclin D1 (among which 5 μM kaempferol can also upregulate the mRNA expression of Pi3k). The 5 μM and 10 μM concentrations of kaempferol have been observed to enhance the phosphorylation of PI3K and AKT1, resulting in elevated p-PI3K/PI3K and p-AKT1/AKT1 ratios. This, in turn, activates the PI3K/AKT signaling pathway, leading to the subsequent phosphorylation of GSK3β. This, in turn, results in the accumulation of β-catenin within the cytoplasm, facilitating its subsequent entry into the nucleus. There, it initiates downstream gene transcription, thereby promoting cell proliferation. A network pharmacology GO analysis reveals that the biological processes primarily involve cell proliferation and its regulation. This finding aligns with the experimental observations made in cell-based studies.

## 6 Conclusion

The present study has demonstrated, through the implementation of cellular experimentation, the capacity of kaempferol to facilitate osteogenic differentiation, in addition to its ability to forestall the incidence and progression of osteoporosis. This finding provides a cogent rationale for the further clinical implementation of kaempferol.

## Availability of Data and Materials

All the data used for the analysis in this study are supported by existing data in the network database, and in this published article, all the data generated in this study are presented.

## Ethics Approval and Consent to Participate

CD and CZ designed the general direction of this study under the guidance of GD. HW and LZ conducted this study. HW wrote the manuscript with the assistance of CZ. WH assisted XY in completing the molecular docking analysis. CZ and DC jointly performed the CCK-8 assay. DG and CZ designed and participated in some experimental content of RT-qPCR and analyzed and calculated the data. XY designed and participated in some experimental contents of the Western Blot and analyzed the experimental data. DC verified the accuracy of all data in the article and provided assistance in the image drawing and layout arrangement. DC and DG corrected language and logical errors in the manuscript. All authors have read and approved the final manuscript. All authors participated sufficiently in the work and agreed to be accountable for all aspects of the work.

## Acknowledgment

Not applicable.

## References

[1] Cosman F, Langdahl B, Leder BZ. Treatment Sequence for Osteoporosis[J]. Endocr Pract, 2024,30(5):490–496. DOI: 10.4158/EP-2023-0727.

[2] Aibar-Almazán A, Voltes-Martínez A, Castellote-Caballero Y, et al. Current Status of the Diagnosis and Management of Osteoporosis [J]. Int J Mol Sci, 2022,23 (16). DOI: 10.3390/ijms23169251.

[3] Chen T, Peng, T. Research progress of Chinese medicine in treating osteoporosis [J]. Journal of Practical Chinese Medicine, 2024, 38(08), 141–143. DOI: 10.13729/j.issn.1671-7813.Z20231934.

[4] Mori T, Komiyama J, Fujii T, et al. Medical expenditures for fragility hip fracture in Japan: a study using the nationwide health insurance claims database [J]. Arch Osteoporos, 2022,17 (1):61. DOI: 10.1007/s11657-022-01100-6.

[5] Moshi M R, Nicopolopoulos K, Stringer D, et al. The Clinical Effectiveness of Denosumab (Prolia®) for the Treatment of Osteoporosis in Postmenopausal Women, Compared to Bisphosphonates, Selective Estrogen Receptor Modulators (SERM), and Placebo: A Systematic Review and Network Meta-Analysis [J]. Calcif Tissue Int, 2023,112 (6):631–646. DOI: 10.1007/s00223-023-01078-3.

[6] Gehrke B, Alves Coelho M C, Brasil D’alva C, et al. Long-term consequences of osteoporosis therapy with bisphosphonates [J]. Arch Endocrinol Metab, 2023,68:e220334. DOI: 10.1590/2359-3997000000542.

[7] Lin Z, Wang S, Liu Z, et al. Exploring Anti-osteoporosis Medicinal Herbs using Cheminformatics and Deep Learning Approaches [J]. Comb Chem High Throughput Screen, 2023,26 (9):1802–1811. DOI: 10.2174/1386207326666230913121117.

[8] Yu Wenlu, Zhang Hong. Research Status of the Action Mechanisms of Chinese Medicinal Materials Containing Active Ingredients in Osteoporosis [J]. China Medicine and Pharmacy, 2024, 14 (21): 32-35+39. DOI: 10.3969/j.issn.2095-0616.2024.21.008.

[9] Wu Peng, Yang Xi, Gao Shang, et al. Mechanism of Herba Cistanches - Epimedium Herb - Fortune’s Drynaria Rhizome in the Treatment of Osteoporosis Based on Network Pharmacology and Molecular Docking [J]. Medical Information, 2024, 37 (13): 1–6. DOI: 10.3969/j.issn.1006-1959.2024.13.001.

[10] Liu H, Yi X, Tu S, et al. Kaempferol promotes BMSC osteogenic differentiation and improves osteoporosis by downregulating miR-10a-3p and upregulating CXCL12 [J]. Mol Cell Endocrinol, 2021,520:111074. DOI: 10.1016/j.mce.2020.111074.

[11] Yang Qipei, Chen Feng, Cui Wei, et al. Signal pathways related to the treatment of osteoporosis with the active monomer kaempferol. China Tissue Engineering Research, 28 (26): 4242–4249. DOI: 10.12102/j.issn.2095-4344.4072.

[12] Brent MB. Pharmaceutical treatment of bone loss: From animal models and drug development to future treatment strategies. Pharmacol Ther. 2023 Apr;244:108383. DOI: 10.1016/j.pharmthera.2023.108383.

[13] Nisha Y, Dubashi B, Bobby Z, Sahoo JP, Kayal S, Ananthakrishnan R, Ganesan P. Cytotoxic Chemotherapy Is Associated with Decreased Bone Mineral Density in Postmenopausal Women with Early and Locally Advanced Breast Cancer. Arch Osteoporos. 2023 Mar 10;18(1):41. DOI: 10.1007/s11657-023-01214-3.

[14] Cosman F, Lewiecki E M, Eastell R, et al. Goal-directed osteoporosis treatment: ASBMR/BHOF task force position statement 2024 [J]. J Bone Miner Res, 2024,39 (10):1393–1405. DOI: 10.1002/jbmr.4963.

[15] Abdi S, Almiman AA, Ansari MGA, Alnaami AM, Mohammed AK, Aljohani NJ, Alenad A, Alghamdi A, Alokail MS, Al-Daghri NM. PTHR1 Genetic Polymorphisms Are Associated with Osteoporosis among Postmenopausal Arab Women. Biomed Res Int. 2021 Dec 22;2021:2993761. DOI: 10.1155/2021/2993761.

[16] Hao Qingfei, Yuan Puwei, Meng Qinghao, et al. Research progress on the mechanism of Icariin in the prevention and treatment of osteoporosis [J]. China Medical Herald, 2024, 21 (10): 189–192. DOI: 10.3969/j.issn.1673-7210.2024.10.041.

[17] Liang Zhou, Zhang Chi, Pan Chengzhen, et al. The mechanism of action of kaempferol in the prevention of osteoporosis based on intestinal flora and extensive targeted metabolomics [J]. Chinese Journal of Tissue Engineering Research, 2025, 29 (20): 4190–4204. DOI: 10.12102/j.issn.2095-4344.4736.

[18] Fan Kaijian, Niu Aiwen, Wu Huihui. Icaritin promotes the proliferation and differentiation of MC3T3-E1 cells through the PI3K/AKT signaling pathway [J]. Zhongnan Pharmacy, 2025, 23 (02): 417–422. DOI: 10.12140/j.issn.1672-2981.2025.02.026.

[19] Yang X, Jiang T, Wang Y, Guo L. The Role and Mechanism of SIRT1 in Resveratrol-regulated Osteoblast Autophagy in Osteoporosis Rats. Sci Rep. 2019 Dec 5;9 (1):18424. DOI: 10.1038/s41598-019-54793-7.

[20] Xi JC, Zang HY, Guo LX, Xue HB, Liu XD, Bai YB, Ma YZ. The PI3K/AKT cell signaling pathway is involved in regulation of osteoporosis. J Recept Signal Transduct Res. 2015;35(6):640–5. DOI: 10.3109/10799893.2015.1070660.

[21] Jing WB, Ji H, Jiang R, Wang J. Astragaloside positively regulated osteogenic differentiation of pre-osteoblast MC3T3-E1 through PI3K/Akt signaling pathway. J Orthop Surg Res. 2021 Oct 7;16(1):579. DOI: 10.1186/s13018-021-02599-3.

[22] Cheng BF, Feng X, Gao YX, Jian SQ, Liu SR, Wang M, Xie YF, Wang L, Feng ZW, Yang HJ. Neural Cell Adhesion Molecule Regulates Osteoblastic Differentiation Through Wnt/ β -Catenin and PI3K-Akt Signaling Pathways in MC3T3-E1 Cells. Front Endocrinol (Lausanne). 2021 May 12;12:657953. DOI: 10.3389/fendo.2021.657953.

[23] Ge L, Cui Y, Liu B, Yin X, Pang J, Han J. ER α and Wnt/ β -catenin signaling pathways are involved in angelicin-dependent promotion of osteogenesis. Mol Med Rep. 2019 May;19(5):3469–3476. DOI: 10.3892/mmr.2019.10124.

[24] Morgan RG, Ridsdale J, Payne M, Heesom KJ, Wilson MC, Davidson A, Greenhough A, Davies S, Williams AC, Blair A, Waterman ML, Tonks A, Darley RL. LEF-1 drives aberrant β -catenin nuclear localization in myeloid leukemia cells. Haematologica. 2019 Jul;104(7):1365–1377. DOI: 10.3324/haematol.2018.205977.

[25] Ni F, Zhang T, Xiao W, Dong H, Gao J, Liu Y, Li J. IL-18-Mediated SLC7A5 Overexpression Enhances Osteogenic Differentiation of Human Bone Marrow Mesenchymal Stem Cells via the c-MYC Pathway. Front Cell Dev Biol. 2021 Dec 17;9:748831. DOI: 10.3389/fcell.2021.748831.

[26] Liu Z, Wu Y. Arctiin elevates osteogenic differentiation of MC3T3-E1 cells by modulating cyclin D1. Bioengineered. 2022 Apr;13(4):10866–10874. DOI: 10.1080/21655979.2022.2064876.

